# Physical activity and screen time in children who survived cancer – A report from the Swiss Childhood Cancer Survivor Study

**DOI:** 10.1101/680140

**Authors:** Christina Schindera, Annette Weiss, Niels Hagenbuch, Maria Otth, Tamara Diesch, Nicolas von der Weid, Claudia E. Kuehni, for the Swiss Pediatric Oncology Group (SPOG) Scientific Committee

**Affiliations:** Swiss Childhood Cancer Registry, Institute of Social and Preventive Medicine, University of Bern, Switzerland; Division of Pediatric Hematology and Oncology, University Children’s Hospital Basel, Switzerland; Department for Epidemiology and Preventive Medicine, Medicine Sociology, University of Regensburg, Germany; Division of Pediatric Hematology and Oncology, University Children’s Hospital Bern, Switzerland

**Author notes:** Corresponding author: Prof. Claudia E. Kuehni, MD; Swiss Childhood Cancer Registry, Institute of Social and Preventive Medicine, University of Bern, Mittelstrasse 43, 3012 Bern, Switzerland. This study has been previously reported as a meeting abstract. Name of presentation: “Physical activity and screen time in children after childhood cancer: A report from the Swiss childhood cancer survivors study,” 2019 National Symposium on Late Complications after Childhood Cancer (NASLCCC), Atlanta, GA, USA, date June 21, 2019, abstract number P48.

**Keywords:** childhood cancer survivors, chronic health conditions, late effects, Swiss Childhood Cancer Registry, exercise

## Abstract

**Background:** Physical activity (PA) can reduce the risk of chronic adverse health conditions in childhood cancer survivors. We examined physical activity and sedentary screen time behavior in a nationwide study in Switzerland.

**Procedures:** The Swiss Childhood Cancer Survivor Study sent questionnaires to parents of all Swiss resident ≥5 year-survivors diagnosed 1995–2010. We assessed physical activity including compulsory school sport, recreational sport, commuting to school, and time spent with screen media in those aged 5–15 years, and compared results to international recommendations.

**Results:** We included 766 survivors with a median age at diagnosis of 2.8 (interquartile range 1.4–5.0) years and a median age at study of 12.5 (10.0–14.3) years. Median PA time was 7.3 (4.8–10.0) hours/week and median screen time 1.4 (0.8–2.0) hours/day. Compulsory school sport hours and walking or cycling to school contributed significantly to total PA. 55% of survivors met PA and 68% screen time recommendations. PA was lower for children living in regions of Switzerland speaking French or Italian compared to German, and for those who had a relapse or musculoskeletal/neurological conditions. Screen time was higher in males, and children with lower parental education and a migration background.

**Conclusions:** PA and sedentary screen watching were associated with social factors and PA also with clinical risk factors. Structural preventions that afford active commuting to school and sufficient school sport are essential, as is counselling vulnerable survivor groups such as those with musculoskeletal and neurological problems, and those who have had a relapse.

## Introduction

Adult childhood cancer survivors have an elevated risk of poor health^1, 2^ and early death^3^: almost 75% suffer from a chronic adverse health condition^2^ and their cumulative mortality reaches nearly 10% 30 years after cancer diagnosis^3^. Physical activity can reduce the risk of cancer and inhibit chronic health conditions like diabetes and hypertension in the general population^4^, while among adult childhood cancer survivors (CCS) physical activity has been associated with reduced risk factors for cardiovascular disease^5^ and cardiovascular disease itself^6^, and with lower overall mortality^7^. Yet while physical activity can mitigate many health hazards^6, 7^, only half of adult CCS meet physical activity recommendations^8-10^.

An active lifestyle might be even more important for young children and teenage survivors, but only a few studies, usually at single centers or with low participant numbers, have been performed in this age group^11-14^. Their results vary, with 31 to 74% meeting recommendations for physical activity^11-15^ and 28 to 46% meeting those for screen time behavior^11, 14^. Research also has neither distinguished between different types of physical activities nor described how school sports or an active daily commute to school might contribute to overall physical activity. Better knowledge of screen time behavior and physical activity and the factors influencing both could inform recommendations for structured prevention and identify risk groups that could profit from counselling or focused interventions.

This nationwide, population-based prospective cohort study in Switzerland investigated activities in the daily life of young CCS and the factors associated with physical activity and screen time.

## Methods

### The Swiss Childhood Cancer Survivor Study

The Swiss Childhood Cancer Survivor Study is a population-based, long-term cohort study of all children registered in the Swiss Childhood Cancer Registry who have been diagnosed since 1976, survived ≥5 years after initial diagnosis, and were alive at the time of the study^16^. The registry includes all patients in Switzerland who were diagnosed at age <21 years with leukemia, lymphoma, central nervous system tumors, malignant solid tumors, or Langerhans cell histiocytosis^17^. Recent estimates indicate that the registry includes 95% of those diagnosed below age 16 since 1995 in Switzerland^18^. We included survivors aged 5–15 years at survey who had been diagnosed between 1995 and 2010. From 2010 to 2016, we traced addresses and sent a questionnaire to parents. We mailed the questionnaire a second time to those who did not respond, and further lack of response included an attempt to reach parents by phone. Among 1068 survivors whose parents were contacted, we received responses from parents of 766 (72%) (Supporting Information TABLE S1, FIGURE S1).

Ethics approval was granted by the Ethics Committee of the Canton of Bern, Switzerland, to the Swiss Childhood Cancer Registry and the Swiss Childhood Cancer Survivor Study (KeK-BE: 166/2014), and the Swiss Childhood Cancer Survivor Study is registered at ClinicalTrials.gov (identifier: NCT03297034).

### Outcomes: physical activity and screen time

We examined physical activity as compulsory school sport, recreational sport, and commuting to school. We derived the time for compulsory school sport, 2.3 hours/week (3 × 45 minutes), from the Swiss school curriculum. Information on recreational sport activities and the commute to school were obtained via questionnaire. Parents were asked about types of recreational sports and how many hours per week CCS devoted to each (Supporting Information FIGURE S2, question 1), and we categorized answers into 16 different types of sports. We also asked how the child usually went to school: on foot or by bike/kickboard, by bus/streetcar, or by car), and the time required (<10 minutes, 10–20 minutes, or >20 minutes); Supporting Information FIGURE S2, questions 2–3); the durations observed in the analysis were 5, 15, and 30 minutes. We considered only transit by foot or bike/kickboard as active. To obtain weekly estimates, we multiplied the times reported for one way to school by 15 to account for five school days per week, and an average of three trips to school per day because there are 2–4 afternoon school sessions per week at Swiss schools, with most children going home for lunch.

We used the World Health Organization (WHO) recommendations to characterize whether a child had sufficient physical activity (≥7 hours/week or ≥60 minutes/day of any physical activity for children aged 5–17 years)^19^. We created the binary outcome of those who had “sufficient” physical activity and those who did not.

Screen time was assessed by asking parents how much time their child spent on average each day interacting with screen media including television, computer games, game boys, PlayStation, or Nintendo (Supporting Information FIGURE S2, question 4). We used the American Academy of Pediatrics (AAP) recommendations for screen-based media exposure to determine acceptable screen time, less than two hours per day^20^. We created the binary outcome of those who had media exposure that was “acceptable” screen time and those who those who did not.

### Clinical characteristics

We extracted the following clinical characteristics from the cancer registry: age at cancer diagnosis, cancer diagnosis, year of cancer diagnosis, treatment protocol, chemotherapy, radiotherapy, surgery, and hematopoietic stem cell transplantation. We classified cancer diagnoses in terms of 12 main groups and Langerhans cell histiocytosis according to the International Classification of Childhood Cancer, third edition (ICCC-3)^21^. We assessed whether children had been treated with anthracyclines. Thoracic radiation included the mantle field, mediastinum, thoracic spine, and total body irradiation (TBI); abdominal radiation included the pelvis, testis, and TBI; and radiation to the head/neck included TBI. We went back to medical records when registry treatment information was incomplete. The questionnaire collected information on chronic health conditions involving the cardiopulmonary and endocrine systems, problems affecting ears and eyes, and musculoskeletal/neurological conditions (Supporting Information TABLE S2).

### Demographic, socioeconomic, and lifestyle characteristics

The questionnaire included demographic (sex, age at study, Swiss language region), socioeconomic (migration background, parental education), and lifestyle characteristics (children’s body mass index [BMI], maternal BMI). We used self-reported weight and heights and calculated children’s BMI and corresponding z-scores^22^. BMI z-scores lower than –2 were classified as underweight, –2 to 1 as normal weight, >1 to 2 as overweight, and >2 as obese ^23^. Self-reported maternal BMI was calculated and categorized according to the National Institutes of Health^24^.

### Statistical analyses

We compared characteristics of participating survivors and those in families from whom we received no response using chi-square tests. We used multivariate imputation by chained equations (MICE) to complete missing values in the outcome variables and demographic, socioeconomic, lifestyle, and clinical variables. Missing values for hours of recreational sport were predicted by corresponding description of recreational sport. All other variables with missing values were imputed by using all other variables with the exception of the outcome variables (Supporting Information text). In an alternative approach, we determined physical activity and screen times using the original data before MICE (Supporting Information TABLE S3). Using multivariable logistic regression, we explored the association between the two binary outcomes, sufficient physical activity (meeting the WHO recommendations) and acceptable screen time (according to AAP recommendations), and demographic, socioeconomic, lifestyle, and clinical characteristics using an a priori selection of clinically important variables. We also investigated the correlation between physical activity and screen time using the pooled Spearman correlation coefficient. We used STATA software (Version 15.1, Stata Corporation, Austin, TX) and R (Version 3.5.2; R Foundation for Statistical Computing, Vienna, Austria)^25^.

## Results

### Study population

The median age at diagnosis of the study population of 766 children (428 were male) was 2.8 years (interquartile range [IQR] 1.4–5.1), median age at survey 12.5 years (10.1–14.3), and median time since diagnosis 9.0 years (7.5–10.8) (TABLE 1). The two most frequent diagnoses were leukemia (37%) and central nervous system tumor (16%), and 51% received anthracyclines and 16% any radiation. Overall, 54% of children reported one or more adverse chronic health condition. At survey, the median BMI z-score in children was 0.08 (–0.7–0.9), and 59% of survivors were normal weight. Full demographic, socioeconomic, and clinical characteristics of CCS are given in Table 1. Participants were comparable to surviving nonparticipants in most characteristics (Supporting Information TABLE S1).

**TABLE 1.**
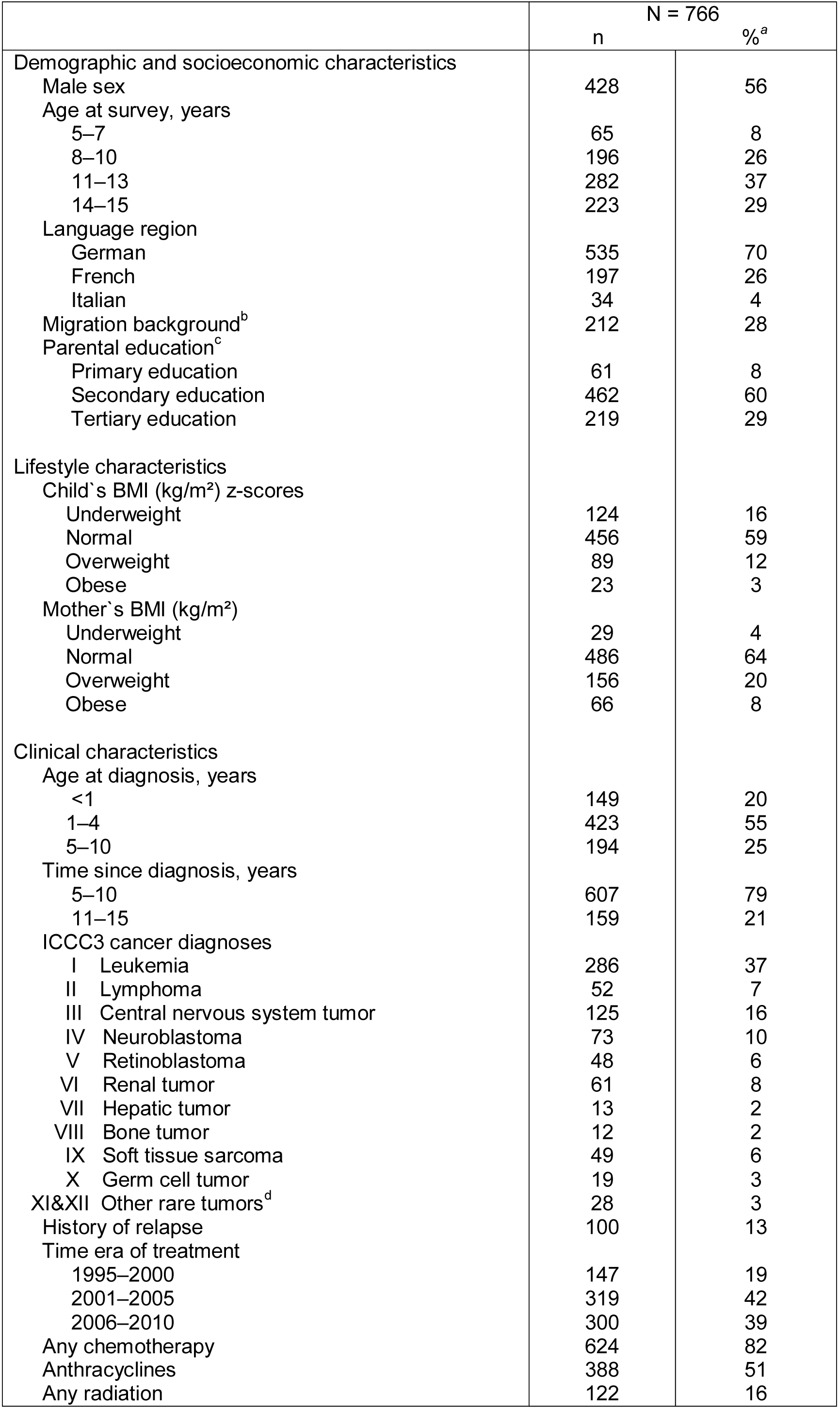
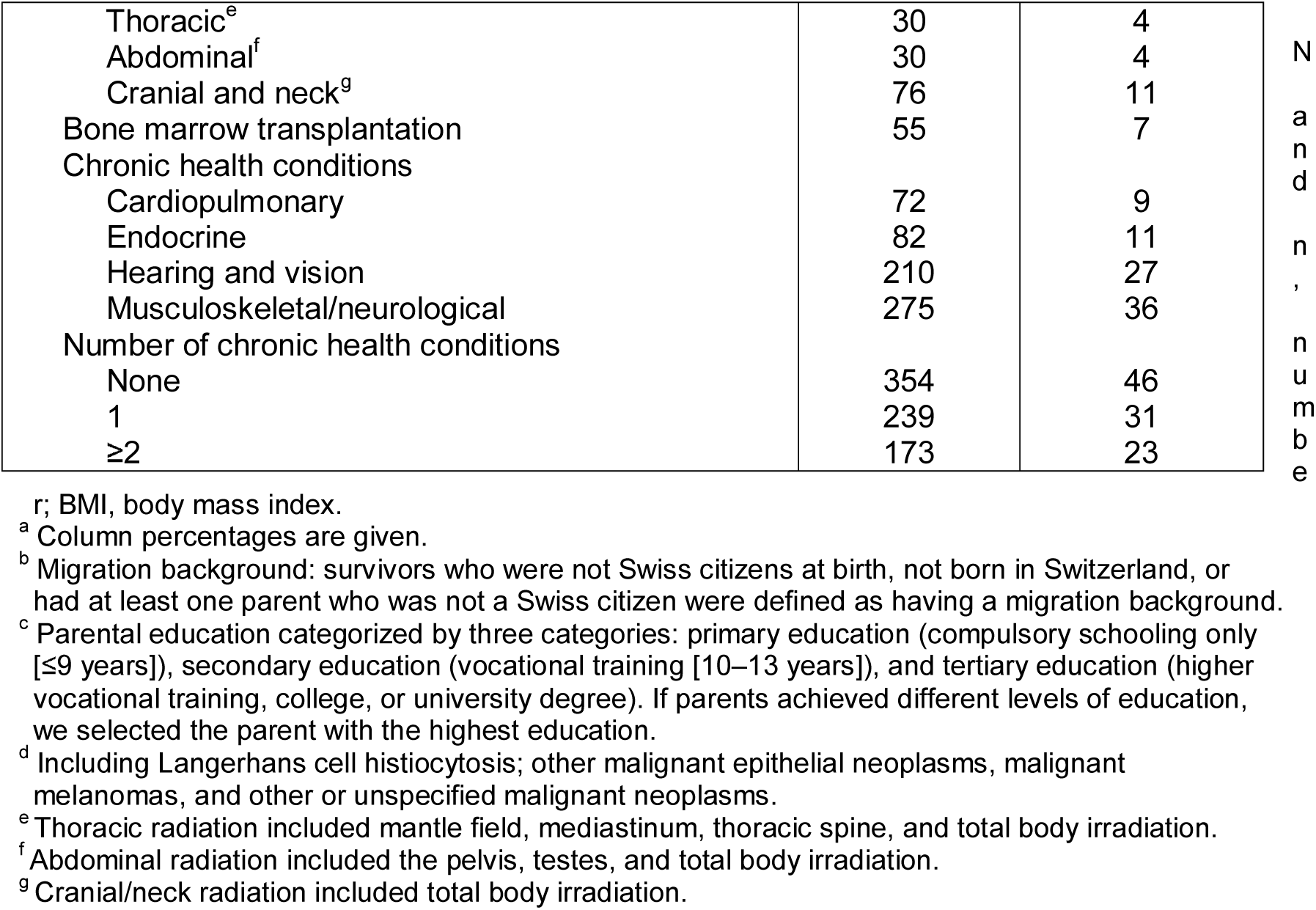
Demographic, socioeconomic, lifestyle, and clinical characteristics of childhood cancer survivors included in the study, N=766

### Physical activity

Overall, the median time devoted to physical activity was 7.3 hours/week, with recreational sport contributing 3.0 hours/week (TABLE 2, FIGURE 1, Supporting Information FIGURE S3). The most common recreational sports were soccer (13%), gymnastics (12%), swimming (11%), cycling/driving a scooter (10%), and free outdoor/indoor play (10%). For male survivors, soccer (21%), scooter (11%), and free indoor/outdoor play (10%) were most relevant, and for female survivors, gymnastics (17%), swimming (13%), and dancing (12%) (FIGURE 2). Over one-half of CCS (55%) had sufficient physical activity according to the WHO recommendations. We found no important difference using the alternative analysis approach that assessed physical activity time and meeting the WHO recommendations using the original data before MICE (Supporting Information TABLE S3).

**TABLE 2.** Compulsory school sport, recreational sport, active way to school, total physical activity, screen time, and adherence to WHO/AAP recommendations in childhood cancer survivors, N=766, 56% males, median age 12.5 years

|  | N=766 |  |  |  |  |
| --- | --- | --- | --- | --- | --- |
|  | Median<sup>a</sup>, IQR | Mean<sup>b</sup>, SD | 10<sup>th</sup>–90<sup>th</sup> percentile | Range | Adherence<sup>c</sup> to WHO/AAP |
| Compulsory school sport (hours/week)<sup>d</sup> | 2.3, NA | 2.3, NA | NA | NA |  |
| Recreational sport (hours/week)<sup>e</sup> | 3.0, 1.0–5.0 | 3.7, 3.9 | 0–8.0 | 0–32 |  |
| Active way to school (hours/week)<sup>f</sup> | 1.3, 0–3.8 | 2.0, 1.9 | 0–3.8 | 0–7.5 |  |
| Total physical activity (hours/week)<sup>g</sup> | 7.3, 4.8–10.0 | 8.0, 4.5 | 3.3–13.5 | 2.3–35.5 | 55% |
| Screen time (minutes/day)<sup>h</sup> | 82, 45–120 | 91, 66 | 15–180 | 0–480 | 68% |

WHO, World Health Organization (recommending $\geq 7$ hours of physical activity per week); AAP, American Academy of Pediatrics, (recommending $< 120$ minutes screen time per day); N, number; IQR, interquartile range; SD, standard deviation; NA, not applicable.
<sup>a</sup> Pooled median over the complete imputed dataset.
<sup>b</sup> Pooled mean over the complete imputed dataset.
<sup>c</sup> Percent of adherence refers to the pooled medians.
<sup>d</sup> Compulsory school sport defined as $3 \times 0.75$ hours/week=2.3 hours/week.
<sup>e</sup> Recreational sport as asked in the questionnaire (question 1, FIGURE S2): Which types of sport does your child perform?
<sup>f</sup> Active way to school by foot or bike/kickboard as asked in the questionnaire (questions 2 and 3, FIGURE S2): How does your child usually go to the kindergarten or to school? How long is your child's way to the kindergarten or school?
<sup>g</sup> Total physical activity consisting of compulsory school sport, recreational sport, and an active way to school. Please indicate, how often your child performs different types of sports (hours per week).
<sup>h</sup> Screen time as asked in the questionnaire (question 4, FIGURE S2): How much time does your child spend on average with the following activities per day? Watching television, computer games, game boy, PlayStation, Nintendo.

**FIGURE 1.**
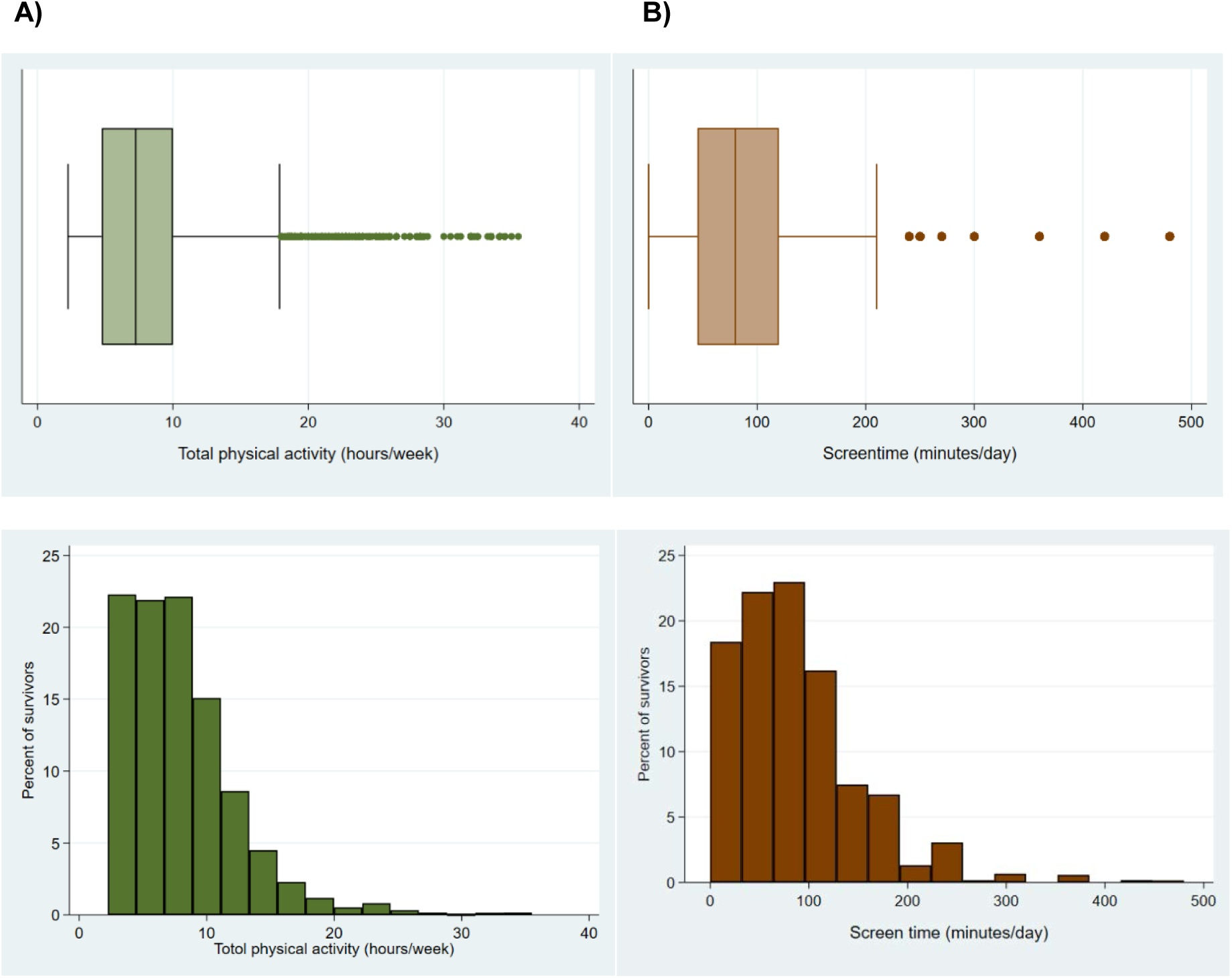
A, box plot and histogram of total physical activity in hours/week; B, box plot and histogram of screen time in minutes/day; reported in childhood cancer survivors (N=766, 56% males, median age 12.5 years)

**FIGURE 2.**
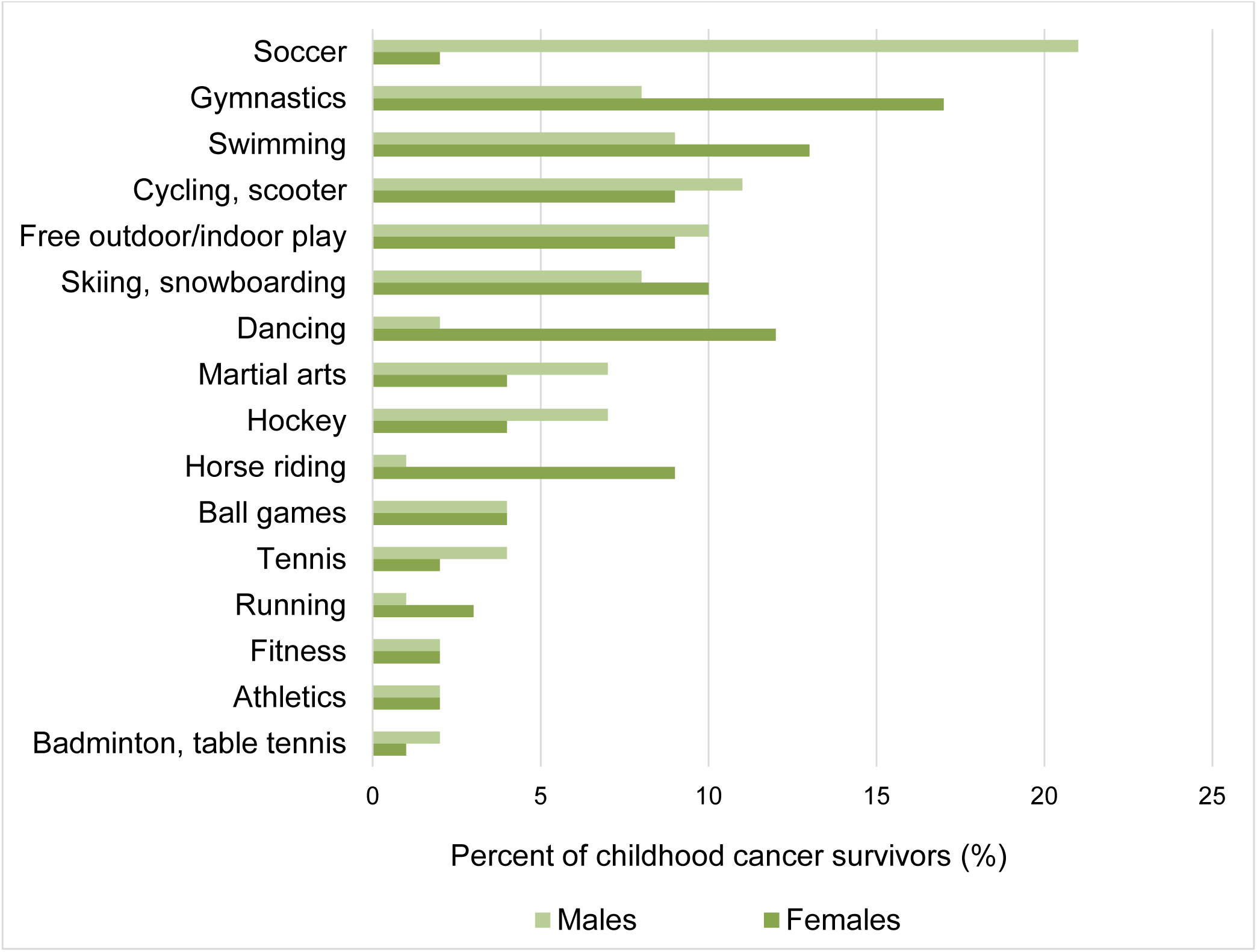
Frequencies of the 16 different recreational sports in childhood cancer survivors (N=766, 56% males, median age 12.5 years), stratified by sex; there can be multiple (1-5) different recreational sports per child

### Screen time

Median screen time was 91 minutes/day, and 68% of children had acceptable screen time in accordance with AAP recommendations (TABLE 2, FIGURE 1). We found no important difference in the alternative analysis approach that assessed screen time and meeting AAP recommendations using the original data before MICE (TABLE S3).

### Predictors for physical activity and screen time

Physical activity was lower for children who lived in the French and Italian language regions than it was in the German-speaking region of Switzerland. It also was lower for those who had had a relapse or suffered from musculoskeletal/neurological conditions (TABLE 3, FIGURE 3). We observed no association between physical activity and sex, age at study, BMI of survivors and mothers, cancer diagnoses, cardiopulmonary conditions, and treatment exposures. Screen time was higher in male survivors and children with lower parental education and migration background (TABLE 3, FIGURE 3), but not associated with sex, age at study, parental education, endocrine and musculoskeletal/neurological problems, and all other clinical characteristics.

**TABLE 3.** Factors associated with a) sufficient physical activity (WHO recommendation) and b) acceptable screen time (AAP recommendation) in childhood cancer survivors, N=766, 56% males, median age 12.5 years

|  | Multivariable logistic regression |  |  |  |  |  |  |  |
| --- | --- | --- | --- | --- | --- | --- | --- | --- |
|  | a) Sufficient physical activity |  |  |  | b) Acceptable screen time |  |  |  |
|  | %<sup>a</sup> | Odds Ratio<sup>b</sup> | 95% CI | p-value<sup>c</sup> | %<sup>a</sup> | Odds Ratio<sup>d</sup> | 95% CI | p-value<sup>c</sup> |
| Demographic characteristics |  |  |  |  |  |  |  |  |
| Sex |  |  |  |  |  |  |  |  |
| Female | 50 | 1.0 |  |  | 72 | 1.0 |  |  |
| Male | 60 | 1.3 | 0.9-1.8 | 0.113 | 64 | 0.6 | 0.4-0.9 | 0.007 |
| Age at study, years |  |  |  |  |  |  |  |  |
| 5–7 | 48 | 1.4 | 0.4–4.4 |  | 88 | 14.5 | 1.1–18.0 | 0.074 |
| 8–10 | 55 | 1.5 | 0.7–3.3 |  | 78 | 2.9 | 1.2–6.9 |  |
| 11–13 | 60 | 1.6 | 1.0–2.6 | 0.306 | 68 | 2.0 | 1.1–3.4 |  |
| 14–15 | 51 | 1.0 |  |  | 52 | 1.0 |  |  |
| Language region |  |  |  |  |  |  |  |  |
| German | 59 | 1.0 |  |  | 68 | 1.0 |  |  |
| French/Italian | 48 | 0.6 | 0.4-0.9 | 0.005 | 65 | 0.8 | 0.6-1.2 | 0.354 |
| Socioeconomic characteristics |  |  |  |  |  |  |  |  |
| Migration background |  |  |  |  |  |  |  |  |
| No | 57 | 1.0 |  |  | 71 | 1.0 |  |  |
| Yes | 51 | 0.9 | 0.6–1.3 | 0.473 | 59 | 0.6 | 0.4–0.9 | 0.018 |
| Parental education |  |  |  |  |  |  |  |  |
| Tertiary education | 60 | 1.0 |  |  | 77 | 1.0 |  |  |
| Secondary education | 54 | 0.8 | 0.6–1.1 | 0.218 | 65 | 0.5 | 0.3–0.8 | 0.004 |
| Primary education | 48 | 0.6 | 0.3–1.1 |  | 50 | 0.4 | 0.2–0.8 |  |
| Lifestyle characteristics |  |  |  |  |  |  |  |  |
| Child's BMI (kg/m<sup>2</sup>) z-scores |  |  |  |  |  |  |  |  |
| Normal weight | 43 | 1.0 |  |  | 66 | 1.0 |  |  |
| Underweight | 50 | 0.8 | 0.5–1.2 | 0.409 | 74 | 1.2 | 0.7–2.1 | 0.803 |
| Overweight/Obesity | 45 | 1.0 | 0.7–1.7 |  | 64 | 0.8 | 0.6–1.6 |  |
| Maternal BMI (kg/m<sup>2</sup>) (continuous) | NA | 1.0 | 0.9–1.0 | 0.381 | NA | 1.0 | 0.9–1.0 | 0.247 |
| Clinical characteristics |  |  |  |  |  |  |  |  |
| Cancer diagnoses |  |  |  |  |  |  |  |  |
| Leukemia | 59 | 1.0 |  |  | 61 | 1.0 |  |  |
| Lymphoma | 60 | 1.1 | 0.6–2.1 | 0.588 | 73 | 1.8 | 0.8–3.8 | 0.111 |
| CNS tumor | 46 | 0.7 | 0.4–1.3 |  | 71 | 2.2 | 1.1–4.7 |  |
| Other tumor | 55 | 0.9 | 0.6–1.4 |  | 71 | 1.3 | 0.8–2.1 |  |
| Relapse |  |  |  |  |  |  |  |  |
| No | 58 | 1.0 |  |  | 67 | 1.0 |  |  |
| Yes | 40 | 0.6 | 0.4–1.0 | 0.030 | 67 | 1.0 | 0.6–1.8 | 0.975 |
| Cardiopulmonary conditions |  |  |  |  |  |  |  |  |
| No | 55 | 1.0 |  |  | 67 | 1.0 |  |  |
| Yes | 56 | 1.1 | 0.7–1.9 | 0.642 | 68 | 1.1 | 0.6–1.9 | 0.805 |
| Endocrine conditions |  |  |  |  |  |  |  |  |
| No | 57 | 1.0 |  |  | 68 | 1.0 |  |  |
| Yes | 42 | 0.6 | 0.4–1.1 | 0.112 | 59 | 0.8 | 0.4–1.4 | 0.360 |
| Hearing/vision chronic conditions |  |  |  |  |  |  |  |  |
| No | 56 | 1.0 |  |  | 66 | 1.0 |  |  |
| Yes | 54 | 1.3 | 0.9–1.9 | 0.195 | 70 | 1.0 | 0.7–1.6 | 0.887 |
| Musculoskeletal/neurological conditions |  |  |  |  |  |  |  |  |
| No | 59 | 1.0 |  |  | 67 | 1.0 |  |  |
| Yes | 49 | 0.7 | 0.5–0.9 | 0.017 | 69 | 1.0 | 0.7–1.5 | 0.936 |
| Anthracyclines |  |  |  |  |  |  |  |  |
| No | 53 | 1.0 |  |  | 68 | 1.0 |  |  |
| Yes | 57 | 0.9 | 0.6–1.3 | 0.506 | 67 | 1.2 | 0.7–1.9 | 0.492 |
| Radiotherapy |  |  |  |  |  |  |  |  |
| No | 55 | 1.0 |  |  | 68 | 1.0 |  |  |
| Yes | 59 | 1.2 | 0.8–1.9 | 0.410 | 66 | 1.1 | 0.7–1.9 | 0.654 |
| Stem cell transplantation |  |  |  |  |  |  |  |  |
| No | 55 | 1.0 |  |  | 68 | 1.0 |  |  |
| Yes | 55 | 1.2 | 0.5–1.5 | 0.543 | 59 | 0.7 | 0.3–1.4 | 0.300 |

WHO, World Health Organization; AAP, American Academy of Pediatrics; N, number; CI, confidence interval; BMI, body mass index; NA, not applicable; CNS, central nervous system
<sup>a</sup> Column percentages are given.
<sup>b</sup> Pooled odds ratios comparing those with high physical activity to those with low physical activity, adjusted for sociodemographic, socioeconomic, and cancer characteristics.
<sup>c</sup> p-value from likelihood ratio tests.
<sup>d</sup> Pooled odds ratios comparing those with low screen time to those with high screen time, adjusted for sociodemographic, socioeconomic, and cancer characteristics.

**FIGURE 3.**
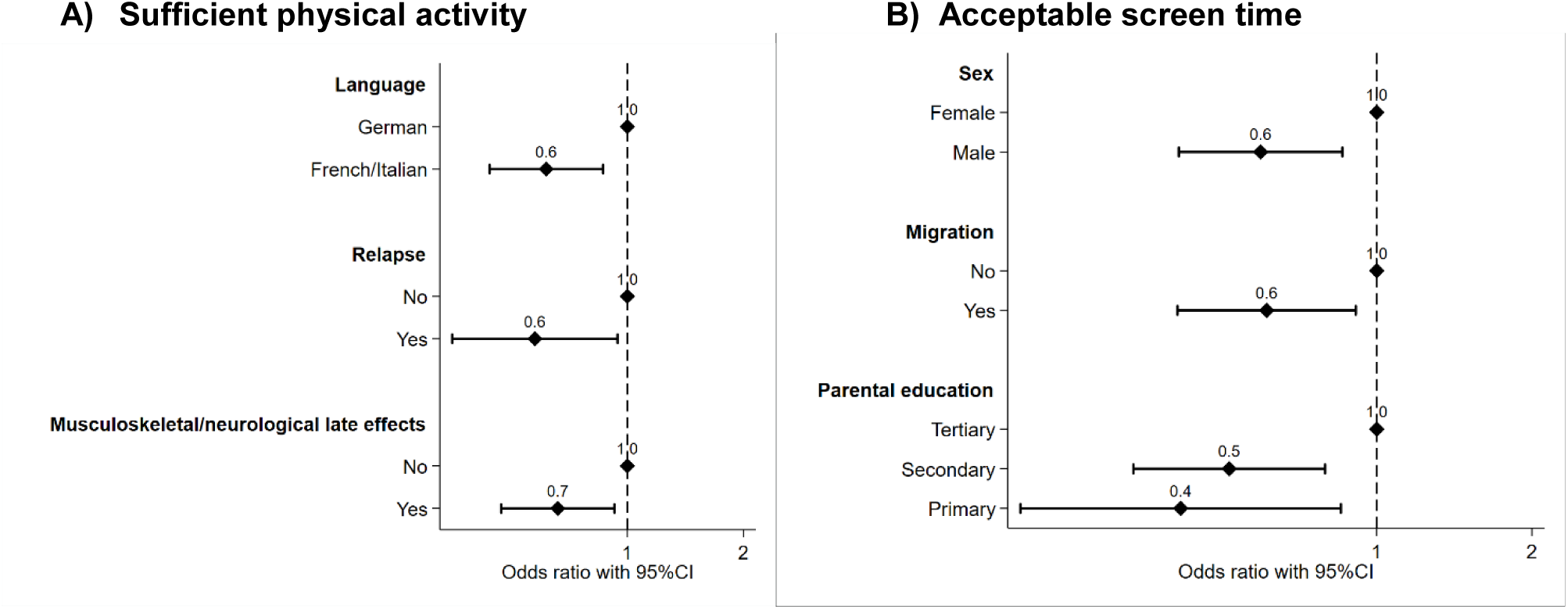
Factors associated with A) sufficient physical activity, and B) acceptable screen time in childhood cancer survivors (N=766, 56% males, median age 12.5 years). Pooled odds ratios from multivariable logistic regression adjusted for demographic, socioeconomic, lifestyle, and clinical characteristics. CI, confidence interval. CNS, central nervous system

### Correlation between low physical activity and high screen times

We found no correlation between physical activity and screen time in survivors (pooled Spearman correlation coefficient –0.05), and no correlation between time spent for recreational sports and an active way to school (pooled Spearman correlation coefficient 0.157) (Supporting Information FIGURE S4).

## Discussion

This comprehensive survey of physical activity and screen time in children and adolescents who have survived cancer found that half of young survivors met the recommendation for physical activity and two-thirds did not exceed the maximum recommended for screen time. Having an active way to get to school and compulsory school sport greatly contributed to overall hours of physical activity.

Our results for physical activity are superior compared to those of a cohort study that included 1300 Swiss children and adolescents between 6–16 years of age, among whom only 39% of children aged 12-13 years met or exceeded physical activity recommendations assessed by accelerometer^26^. Also in that study, children who lived in the French and Italian linguistic regions of Switzerland were less active than those living in the German-speaking region^26^. Young survivors in our study prefered recreational sports including soccer for males, and gymnastics, swimming, and dancing for females, and these preferences are similar to those of other school children^27^. A study of screen time in healthy adolescent school children in Switzerland found higher median screen times of 122 minutes per day (compared to 82 minutes per day in our population), though screen time was assessed differently and focused on internet use^28^.

Studies of physical activity and screen time in children after cancer are few and report variable results. A 2012 Australian study assessed 40 children in two centers after hematopoietic stem cell transplantation at a mean age of 12.5 years by questionnaire; 48% of the children met the physical activity recommendation and 28% that for screen time^11^. Another single-center Australian study that used a three-day diary assessed 74 young survivors with a mean age of 15.0 years between 2012 and 2014 and reported that 74% adhered to physical activity and 46% to screen time recommendations^14^. Gilliam and colleagues performed phone interviews between 2010 and 2011 in 105 North American survivors aged 11.1 years who reported a mean physical activity time of 6.7 hours per week, which is lower than the mean 8.0 hours we observed^12^, but times since diagnosis differed between the two cohorts (9.0 years in our study versus 4.6 years in the North American study). Another study from two North American centers reported a mean physical activity time of 47 minutes per day for 319 survivors aged 14.6 years, which corresponds to 5.5 hours per week and again is lower than in our cohort^15^. But that study used questionnaires that focused on past-year leisure-time physical activity, whereas our questionnaire also included the way to school and school sport. This could explain the difference.

Important predictors for physical activity in young survivors in other studies are social support from family and peers^12^. Additional predictors reported for adolescent and adult survivors include female sex^8, 9^, low parental^8^ or survivor education^9, 13, 29^, cranial radiation^8, 9, 29^, overweight and obesity^9, 29^, physical limitations^8, 29^, and a diagnosis of central nervous system tumors and sarcomas^29, 30^. We also found that musculoskeletal/neurological problems are a predictor for physical activity and we also saw a trend for parental education, central nervous system tumors, and endocrine conditions.

Among our study’s limitations is the reliance of the outcome variables physical activity and screen time on parental reporting. Parents might have overestimated physical activity and underestimated screen time because of both social desirability and recall biases. A second problem involves the questionnaire’s having inquired about structured physical activities even though young children in particular are active mainly in an unstructured way with free inside and outside play^26^. This differential misclassification bias could have led to underestimation of activity in younger survivors. However, parents did mention free outdoor and indoor play in 10% of boys and 9% of girls (FIGURE 2). Accelerometers and pedometers may overcome this problem^26, 31^, though worn only for study purposes and short periods their data might not be representative of daily life. A third limitation is that physical activity in Swiss school children differs between winter and summer^26^, but we did not account for the season when parents filled out the questionnaire. Fourth, screen time questions assessed traditional screen activities such as watching television and computer games, and not use of social media and mobile phones; the average screen time we observed might be an underestimate. Finally, our study had no control group because too few siblings met our inclusion criteria.

This study is the first nationwide, population-based study of physical activity and screen time in children who survived cancer. Among its strengths is its relatively high response rate, 72%, which makes us confident that the results are representative for Swiss CCS. Also, being nested in the Swiss Childhood Cancer Registry provided us with important, comprehensive data on demographic, socioeconomic, lifestyle, and clinical characteristics.

Our results indicate that structural support via compulsory school sport and an active daily commute to school are important contributors to physical activity. Public health policy should at least preserve if not increase support for both compulsory and voluntary school sport. Further research should inquire into why physical activity is lower in the parts of Switzerland speaking French and Italian than in the German-speaking part, and how physical activity levels might be increased in all three. Also, it goes without saying that family and community support for actively commuting to school should be maintained or increased.

For individual prevention, clinicians should counsel young survivors and their families to pursue active lives. In a German study, only 25% of 83 young survivors with a median age of 14 years and 3.8 years after cancer diagnosis participated in school sport, and medical advisories against sports participation were frequent^32^. Parents also might overprotect their children during and after completion of cancer therapy. Physical activity not only is safe both during and after cancer therapy, it may positively influence evolving chronic health conditions. Pediatric oncologists therefore can and should assure families that physical activity is of particular importance to CCS and encourage participation in compulsory and voluntary school sports, and keep medical restrictions on activity to a minimum. Further research should include interventions that include social support as an important contributor to children’s physical activity^12^.

In summary, we found that half of young cancer survivors are not active enough and one-third devote too much time to sedentary screen viewing. Compulsory school sport and an active commute to school are important components of an active lifestyle. Therefore, we need both individual-based prevention such as better counselling of survivors and families, and structural prevention addressed to all children in Switzerland, such as promotion of active commuting to school and extended school sport lessons.

## Abbreviation table

AAP: American Academy of Pediatrics
BMI: Body mass index
CCS: Childhood cancer survivor
CI: Confidence Interval
ICCC-3: International Classification of Childhood Cancer, third edition
IQR: Interquartile range
MICE: Multivariate imputation by chained equations
OR: Odds ratio
TBI: Total body irradiation
WHO: World Health Organization

## Acknowledgements

We thank all childhood cancer survivors and families for participating in our survey. We thank the study team of the SCCSS (Fabiën Belle, Carole Dupont, Rahel Kasteler, Rahel Kuonen, Nadine Lötscher, Jana Remlinger, Grit Sommer, Nicolas Waespe, and Annette Weiss,) the data managers of the SPOG (Dr. Claudia Althaus, Nadine Assbichler, Pamela Balestra, Heike Baumeler, Nadine Beusch, Sarah Blanc, Dr. Pierluigi Brazzola, Susann Drerup, Janine Garibay, Franziska Hochreutener, Monika Imbach, Friedgard Julmy, Eléna Lemmel, Rodolfo Lo Piccolo, Heike Markiewicz, Dr. Veneranda Mattielo, Annette Reinberg, Dr. Renate Siegenthaler, Astrid Schiltknecht, Beate Schwenke, and Verena Stahel) and the team of the SCCR (Meltem Altun, Erika Brantschen, Katharina Flandera, Elisabeth Kiraly, Verena Pfeiffer, Shelagh Redmond, Julia Ruppel, Ursina Roder). We thank Christopher Ritter for editorial assistance.

This study was supported by the Swiss Cancer League (grant numbers: KLS-3886-02-2016, KFS-4157-02-2017). The work of the Swiss Childhood Cancer Registry is supported by the Swiss Pediatric Oncology Group (www.spog.ch), Schweizerische Konferenz der kantonalen Gesundheitsdirektorinnen und –direktoren (www.gdk-cds.ch), Swiss Cancer Research (www.krebsforschung.ch), Kinderkrebshilfe Schweiz (www.kinderkrebshilfe.ch), the Federal Office of Public Health (FOPH) and the National Institute of Cancer Epidemiology and Registration (www.nicer.org).

## Data availability statement

The Swiss Childhood Cancer Registry and Swiss Childhood Cancer Survivor Study are a collaborative project of the Swiss Pediatric Oncology Group (SPOG) and the Institute of Social and Preventive Medicine, University of Bern, Switzerland. Our homepage displays detailed information in methods, results and publications (www.childhoodcancerregistry.ch). Researchers interested in collaborative work can contact the corresponding author (Claudia Kuehni) to discuss planned projects or analyses of existing data. The final decision will be made upon presentation of the project to the Scientific Council of the Swiss Pediatric Oncology Group^16^.

## Supporting Information

**TABLE S1** Demographic, socioeconomic, and clinical characteristics of responding and nonresponding childhood cancer survivors (N=1068)

**TABLE S2** Selected chronic health conditions asked in the Swiss Childhood Cancer Survivor Study for survivors aged 5–15 years and included in this study

**TABLE S3** Compulsory school sport, recreational sport, active way to school, total physical activity, screen time, and adherence to WHO and AAP recommendations in childhood cancer survivors, N=766, 56% males, median age 12.5 years; original data before multivariate imputation by chained equations (MICE)

**FIGURE S1** Population tree of Swiss childhood cancer survivors eligible for the study, contacted and responding to the questionnaire of the Swiss Childhood Cancer Survivor Study

**FIGURE S2** Physical activity and screen time, asked in the Swiss Childhood Cancer Survivor Study questionnaire: recreational sport (question 1), the way to school (questions 2 and 3), and screen time (question 4)

**FIGURE S3** Pooled means of recreational sport, compulsory school sport, and active way to school by foot, bike/kickboard contributing to the total physical activity in hours/week in childhood cancer survivors (N=766, 56% males, median age at study 12.5 years)

**FIGURE S4** Scatterplots of A) total physical activity and screen time, and B) recreational sport and active way to school, evaluated in childhood cancer survivors, N=766, 56% males, median age at study 12.5 years; no correlation between variables (pooled Spearman correlation coefficient for total physical activity and screen time −0.05; pooled Spearman correlation coefficient for recreational sport and active way to school 0.157)

**Supporting information text S1**: Description of number of missing values

