## Supporting Information for "Physical activity and screen time in children who survived cancer – A report from the Swiss Childhood Cancer Survivor Study"

**TABLE S1** Demographic, socioeconomic, and clinical characteristics of responding and nonresponding childhood cancer survivors (N=1068)

|  | Responder, n=766  n %*^a^* | | Nonresponder, n=302  n %*^a^* | | *P^b^* |
| --- | --- | --- | --- | --- | --- |
| Demographic characteristics  Male sex  Age at survey, years  Median [IQR]  5–7  8–10  11–13  14–15  Language region  German  French  Italian | 428  12.5 [10.0–14.3]  65  196  282  223  535  197  34 | 56  8  26  37  29  70  26  4 | 166  12.1 [9.9–14.3]  26  76  112  88  207  83  12 | 55  9  25  37  29  69  27  4 | *0.788*  *0.960*  *0.811* |
| Socioeconomic characteristics  Migration background^c^ | 212 | 28 | 147 | 49 | *<0.001* |
| Clinical characteristics  Age at diagnosis, years  Median [IQR]  <1  1–4  5–10  Time since diagnosis, years  Median [IQR]  5–10  11–15  ICCC3 cancer diagnoses  I Leukemia  II Lymphoma  III CNS tumor  IV Neuroblastoma  V Retinoblastoma  VI Renal tumor  VII Hepatic tumor  VIII Bone tumor  IX Soft tissue sarcoma  X Germ cell tumor  XI & XII Other rare tumors^d^  History of relapse  Time era of treatment  1995–2000  2001–2005  2006–2010  Any chemotherapy | 2.8 [1.4–5.1]  149  423  194  8.5 [7.0–10.2]  607  159  286  52  125  73  48  61  13  12  49  19  28  100  147  319  300  624 | 20  55  25  79  21  37  7  16  10  6  8  2  2  6  3  3  13  19  42  39  82 | 2.9 [1.4–4.8]  57  175  70  8.5 [6.8–10.2]  238  64  101  27  60  30  13  23  6  6  12  9  15  45  55  114  133  232 | 19  58  23  79  21  33  9  20  10  4  8  2  2  4  3  5  15  18  38  44  77 | *0.695*  *0.875*  *0.571*  *0.428*  *0.334*  *0.076* |

N, number; IQR, interquartile range; CNS, central nervous system

^a^ Column percentages are given.

^b^ p-values calculated from chi-squared tests comparing responders and non-responders.

^c^ Migration background: survivors who were not Swiss citizens at birth, not born in Switzerland, or had at least one parent who was not a Swiss citizen were defined as having a migration background.

^d^ Including Langerhans cell histiocytosis; other malignant epithelial neoplasms, malignant melanomas, and other or unspecified malignant neoplasms.

**SUPPORTING INFORMATION**

**TABLE S2** Selected chronic health conditions asked in the Swiss Childhood Cancer Survivor Study for survivors aged 5–15 years and included in this study

| Cardiopulmonary | Endocrine | Hearing and vision | Musculoskeletal/  neurological |
| --- | --- | --- | --- |
| Cardiac   - Arrhythmia^1^ - Cardiomyopathy - Myocardial infarction - Pericarditis - Valvular problems - Stroke - Deep vein thrombosis or pulmonary embolism   Pulmonary   - Chronic cough^2^ - Recurrent pneumonias - Lung fibrosis - Emphysema - Chest wall abnormalities | - Hypothyroidism - Hyperthyroidism - Diabetes^3^ - Diabetes^4^ - Diabetes insipidus^5^ - Growth hormone deficiency | Hearing   - Hearing loss^6^ - Deafness - Tinnitus^7^   Vision   - Moderate visual impairment - Blindness^8^ - Cataract - Very dry eyes^9^ - Double vision - Eye movement disorders^10^ | Musculoskeletal   - Shortened extremities - Reduced flexibility of joints - Prolonged pain in bones or joints - Scoliosis   Neurological   - Weakness or inability to move arms or legs - Decreased sense of touch or feeling - Problems with balance - Problems chewing or swallowing - Loss of taste or smell - Speech^11^ - Epilepsy^12^ |

^1^ Requiring follow-up by a doctor

^2^ For greater than three months

^3^ Controlled with diet or tablets

^4^ Controlled with insulin shots

^5^ Controlled with Minirin (Desmopressin)

^6^ Not completely corrected by hearing aid

^7^ Or ringing in the ears

^8^ In one or both eyes

^9^ Requireing eye drops

^10^ Including ocular muscle palsy

^11^ Including stammering or stuttering

^12^ Including convulsions and blackouts

**SUPPORTING INFORMATION**

**TABLE S3** Compulsory school sport, recreational sport, active way to school, total physical activity, screen time, and adherence to WHO and AAP recommendations in childhood cancer survivors, N=766, 56% males, median age 12.5 years; original data before multivariate imputation by chained equations (MICE)

|  |  | N=766 | | | | |
| --- | --- | --- | --- | --- | --- | --- |
|  | Median, IQR | | Mean, SD | 10^th^–90^th^ percentile | Range | Adherence to WHO/AAP |
| Compulsory school sport (hours/week)^a^  Recreational sport (hours/week)^b^  Active way to school (hours/week)^c^ | 2.3, NA  2.5, 1.0–4.5  1.3, 1.3–3.8 | | 2.3, NA  3.4, 3.8  2.1, 1.9 | NA  0–7.5  0–3.8 | NA  0–30.0  0–7.5 |  |
| Total physical activity (hours/week)^d^ | 7.0, 3.9–9.5 | | 7.4, 4.5 | 2.3–13.0 | 2.3–34.1 | 51% |
| Screen time (minutes/day)^e^ | 80, 45–120 | | 91, 66 | 15–180 | 0–480 | 60% |

WHO, World Health Organization (recommending ≥7 hours of physical activity per week); AAP, American Academy of Pediatrics, (recommending <120 minutes screen time per day); N, number; IQR, interquartile range; SD, standard deviation; NA, not applicable.

^a^ Compulsory school sport defined as 3 x 0.75 hours/week=2.3 hours/week.

^b^ Recreational sport as asked in the questionnaire (question 1, FIGURE S2): Which types of sport does your child perform?

^c^ Active way to school by foot or bike/kickboard as asked in the questionnaire (questions 2–3, FIGURE S2): How does your child usually go to the kindergarten or to school? How long is your child’s way to the kindergarten or school?

^d^ Total physical activity consisting of compulsory school sport, recreational sport, and an active way to school. Please indicate, how often your child performs different types of sports (hours per week).

^e^ Screen time as asked in the questionnaire (question 4, FIGURE S2): How much time does your child spend on average with the following activities per day? Watching television, computer games, game boy, play station, Nintendo.

**SUPPORTING INFORMATION**

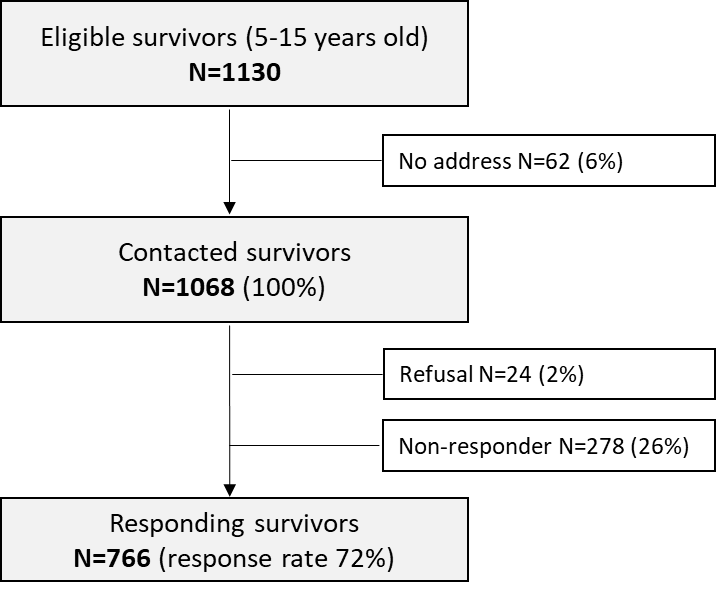

**FIGURE S1** Population tree of Swiss childhood cancer survivors eligible for the study, contacted and responding to the questionnaire of the Swiss Childhood Cancer Survivor Study

**SUPPORTING INFORMATION**

| 1. | Which types of sport does your child perform? Please indicate, how often your child performs different types of sport. | | | | | | | | | | | |
| --- | --- | --- | --- | --- | --- | --- | --- | --- | --- | --- | --- | --- |
|  | Types of sport | | | | | | | |  | | | Hours per week |
| 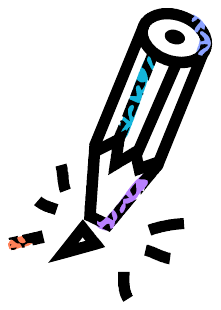 |  | | | | | | | |  | | |  |
| 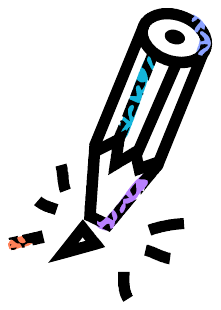 |  | | | | | | | |  | | |  |
| 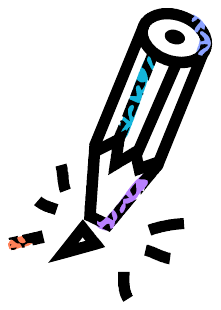 |  | | | | | | | |  | | |  |
| 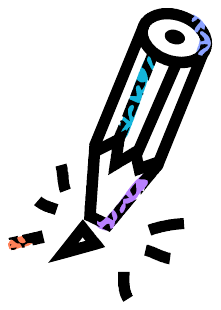 |  | | | | | | | |  | | |  |
| 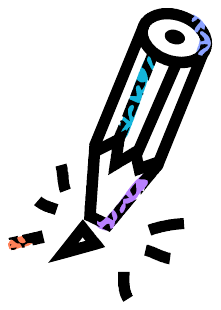 |  | | | | | | | |  | | |  |

| 2. | How does your child usually go to the kindergarten or to school? | | | |
| --- | --- | --- | --- | --- |
|  | ☐ | By foot | | |
|  | ☐ | By bike/kickboard | | |
|  | ☐ | By bus/streetcar | | |
|  | ☐ | 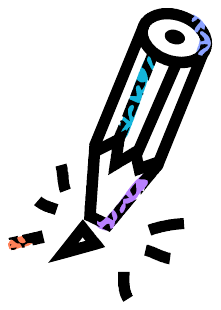By car | | |
|  | ☐ | By the following: | | _____________________ |
| 3. | How long is your child`s way to the kindergarten or school (only one way)? | | | |
|  | ☐ | Less than 10 minutes | | |
|  | ☐ | 10 – 20 minutes | | |
|  | ☐ | More than 20 minutes | | |

| 4. | How much time does your child spend on average with the following activities per day? | | | | | |
| --- | --- | --- | --- | --- | --- | --- |
|  | Watching television: | | | | ___________ | minutes/day |
|  | Computer games, game boy, play  station, Nintendo: | | | | ___________ | minutes/day |

**FIGURE S2** Physical activity and screen time, asked in the Swiss Childhood Cancer Survivor Study questionnaire: recreational sport (question 1), the way to school (questions 2 and 3), and screen time (question 4)

**SUPPORTING INFORMATION**

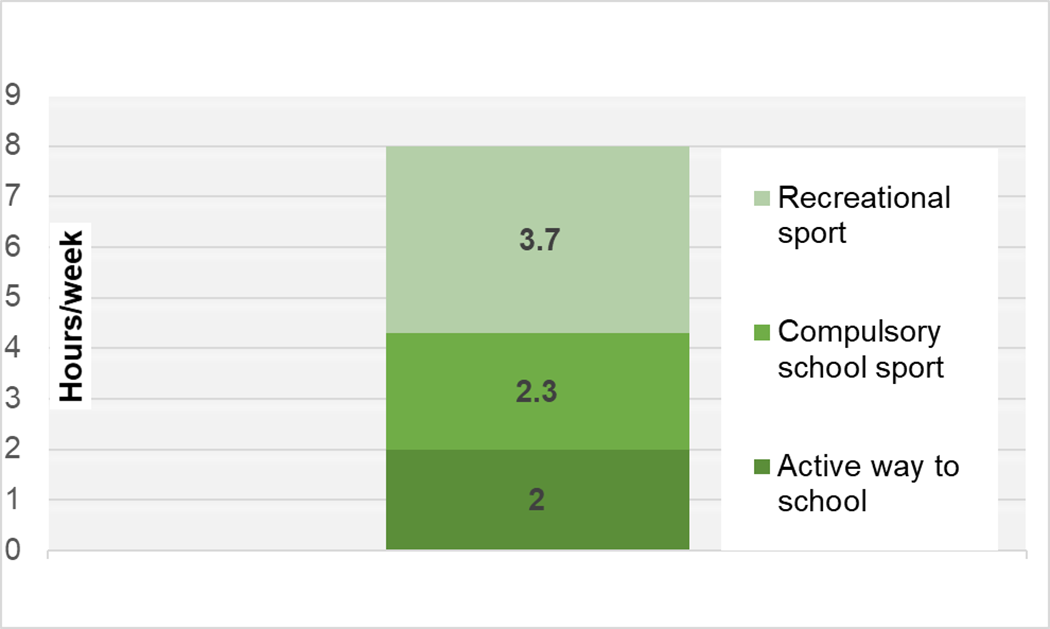

**>50%**

**FIGURE S3** Pooled means of recreational sport, compulsory school sport, and active way to school by foot, bike/kickboard contributing to the total physical activity in hours/week in childhood cancer survivors (N=766, 56% males, median age at study 12.5 years)

**SUPPORTING INFORMATION**

A) B)

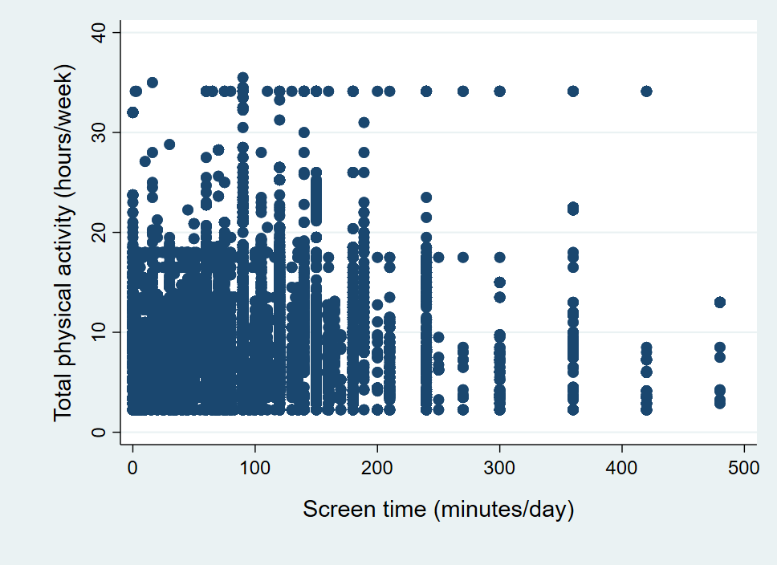

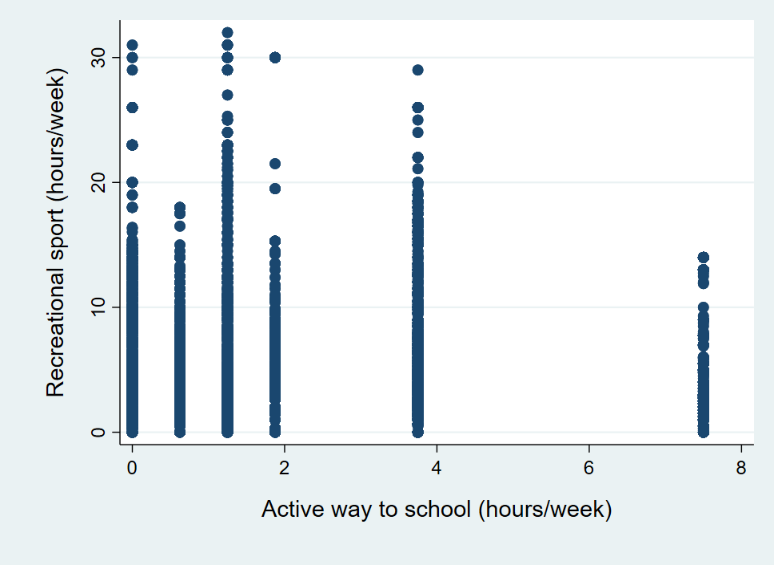

**FIGURE S4** Scatterplots of A) total physical activity and screen time, and B) recreational sport and active way to school, evaluated in childhood cancer survivors, N=766, 56% males, median age at study 12.5 years; no correlation between variables (pooled Spearman correlation coefficient for total physical activity and screen time -0.05; pooled Spearman correlation coefficient for recreational sport and active way to school 0.157)

**SUPPORTING INFORMATION**

**TEXT** Description of number of missing values

Organ-specific late effects were not always indicated. With the assumption that such grave health issues would be known and stated, missing values were set to “absent.” As a consequence, these variables (and their sum) are free of missing values. For the eight children who did not state whether they are doing sports or not, the same approach was taken.

On the observational level, 79% of the cases were complete, the rest were missing data in up to three variables simultaneously. Missing values were present in six of the 18 a priori selected covariates (BMI of the child, BMI of the mother, parental education, anthracycline treatment, hematopoietic stem cell transplantation, and radiotherapy). For the total physical activity outcome, the (up to five) physical activities, school sport, and way to school showed varying numbers of missing values: of the 567 indications of a first physical activity, 31 (5.5%) lacked the time; of the 364 indicated second activities, 43 (11.8%) had no time mentioned; of 165 third physical activities, 29 (17.6%) missed the time; of the 59 fourth physical activities, 19 (32.2%) had no time; and of the 19 fifth physical activities, 3 (15.8%) did not state the time. The time to school was missing in 98 instances (12.8%). The outcome screen time was absent in 90 cases (11.7%).

To avoid loss of power and severe underestimation of total physical activity and screen time, the missing values were imputed using multivariate imputation by chained equations (R package mice, version 3.3.0) (1). The assumptions for the imputation of the physical activity time were that a lack of a description of physical activity indicated no physical activity (0 mins. per week), and that an absent time with a description provided was missing completely at random (MCAR). The missing information in the covariates was assumed to be missing at random (MAR).

A missing description of activity was never imputed. Each missing activity time was only predicted by its corresponding description (if present). Activity descriptions and times were not used to predict missing values in other covariates. The total sum of activities was calculated passively after its summands (up to five activity times, school sport, and way to school) were completed. The other covariates with missing values were imputed by using all other variables (with the exception mentioned above), thus addressing the assumption of MAR. No automatic variable predictor was used.

As imputation method, categorical variables were imputed with the logistic or mulitnomial logit model, while for all continuous variables, predictive mean matching was used. To minimize simulation error, 100 imputed data sets were generated with 50 iterations per imputation to ascertain convergence. The results were inspected visually and showed good convergence, absence of patterns in the imputed data sets, and meaningful imputed values.

The analyses were performed on each of the 100 imputed data sets, and the results were pooled according to Rubin’s rules (2). As diagnostics to assess how strongly the estimated parameters may be influenced by missing data, the fraction of missing information due to nonresponse (FMI) and the proportion of the total variance attributable to the missing data (Lambda) are given.
